# Novel and known signals of selection for fat deposition in domestic sheep breeds from Africa and Eurasia

**DOI:** 10.1101/493940

**Authors:** Salvatore Mastrangelo, Hussain Bahbahani, Bianca Moioli, Abulgasim Ahbara, Mohammed Al Abri, Faisal Almathen, Anne da Silva, Baldassare Portolano, Joram M. Mwacharo, Olivier Hanotte, Fabio Pilla, Elena Ciani

**Author notes:** These authors contributed equally to this work (Hussain Bahbahani, Bianca Moioli).

## Abstract

Genomic regions subjected to selection frequently show signatures such as within-population reduced nucleotide diversity and outlier values of differentiation among differentially selected populations. In this study, we analyzed 50K SNP genotype data of 373 animals belonging to 23 sheep breeds of different geographic origins using the *Rsb* and *F_ST_* statistical approaches, to identify loci associated with the fat-tail phenotype. We also checked if these putative selection signatures overlapped with regions of high-homozygosity (ROH). The analyses identified novel signals and confirmed the presence of selection signature in genomic regions that harbor candidate genes known to affect fat deposition. Several genomic regions that frequently appeared in ROH were also identified within each breed, but only two ROH islands overlapped with the putative selection signatures. The results reported herein provide the most complete genome-wide study of selection signatures for fat-tail in African and Eurasian sheep breeds; they also contribute insights into the genetic basis for the fat tail phenotype in sheep, and confirm the great complexity of the mechanisms that underlie quantitative traits, such as the fat-tail.

## Introduction

Natural selection plays an important role in determining the individuals that are best adapted to novel and existing environmental conditions. Besides natural selection, artificial selection has been widely applied to livestock species to achieve more desirable/profitable phenotypes [1]. For instance, sheep *(Ovis aries)* have been selected since domestication, approximately 9,000 years ago [2]. This process of selection resulted in divergent sheep breeds, reared in different geographic regions due to their different adaptability. Among these, fat-tail are an important class of sheep breeds and represent about 25% of the world’s sheep population [3] mainly distributed in the Middle East, North and East Africa and Central Asia. According to Xu et al. [4] fat tails represent the energy reserve necessary to survive critical conditions such as drought seasons and food shortage. This statement being emphasized by Mwacharo et al. [5] who confirmed that the fat-tails are the predominant sheep across the deserts of northern Africa, and in the highlands, semi-arid and arid environments of eastern and southern Africa while the thin-tails occur in Sudan and in the sub-humid and humid regions of West Africa.

The unique genetic patterns inscribed in the genome of individuals by natural and/or artificial selection are defined as signatures of selection, which are usually regions of the genome that harbor functionally important sequence variants [6]. Although human consumption of animal fat has dramatically reduced in preference of leaner meat, the investigation of the potential candidate genes involved in the fat-tail might contribute to exploring the genetics of fat deposition, energy storage and adaptation to climate changes [7–9]. With the aim to identify candidate genes with a potential role in these traits, several authors performed studies targeting the fat-tail phenotype contrasted with the thin-tail one. All authors used, for their comparisons, sheep of the same geographic regions to prevent referring to the fat-tail differentiation signals arising from different origins or isolation by distance. These studies included indigenous Chinese [4,8,10,11], Mediterranean-North African [9,12], Iranian [3] and Ethiopian breeds [13]. Similarly, in this study, the genomes of fat-tail sheep from different regions of Africa and Eurasia were contrasted with the genomes of thin-tail sheep of the same geographical area. In order to improve the specificity of signal detection, we combined two complimentary approaches (*Rsb* and *F_ST_*); moreover, as distinguishing false positive genes from candidate genes is not straightforward, selection signatures of fat-tail were considered only when shared by two or more fat-tail breeds of different geographic origin, and substantiated by verifying whether the candidate genes in proximity of the anonymous markers of differentiation have a known, or assumed, role in fat deposition and adipogenesis in mammals.

## Materials and Methods

### Samples, genotyping and quality control

A total of 373 animals belonging to 23 sheep breeds from different geographic regions were selected (Table 1). For all the animals, genotype data from the Illumina OvineSNP50 BeadChip array were collated for the analysis. Chromosomal coordinates for each SNP were obtained from the latest release of the ovine genome sequence assembly Oar_v4.0. The dataset was filtered to remove animals with more than 10% missing genotypes, SNPs with a call rate lower than 95% and with a minor allele frequency (MAF) lower than 1%, and to exclude non-autosomal and unassigned markers.

**Table 1.** Breeds, number of animals (N), tail type, country and origin of genotyping data of the breeds used in the contrasting groups (fat-vs. thin-tail).

| Breed | N | Tail type | Country | Comparison | Data origin |
| --- | --- | --- | --- | --- | --- |
| Bonga | 9 | long fat-tail | Ethiopia | 1, 2 | [13] |
| Doyogena | 15 | long fat-tail | Ethiopia | 1, 2 | [13] |
| Gesses | 10 | long fat-tail | Ethiopia | 1, 2 | [13] |
| Kido | 10 | long fat-tail | Ethiopia | 1, 2 | [13] |
| Loya | 15 | long fat-tail | Ethiopia | 1, 2 | [13] |
| Shubi Gemo | 15 | long fat-tail | Ethiopia | 1, 2 | [13] |
| Kefis | 14 | rumped | Ethiopia | 1 | [13] |
| Arabo | 10 | rumped | Ethiopia | 1 | [13] |
| Adane | 12 | rumped | Ethiopia | 1 | [13] |
| Molale (Menz) | 15 | short fat-tail | Ethiopia | 1 | [13] |
| Gafera (Washera) | 15 | short fat-tail | Ethiopia | 1 | [13] |
| Huri | 7 | fat-tail | Arabian peninsula | 3 | [13] |
| Naimi | 7 | fat-tail | Arabian peninsula | 3 | [13] |
| Najdi | 6 | fat-tail | Arabian peninsula | 3 | [13] |
| Omani | 10 | fat-tail | Arabian peninsula | 3 | [13] |
| Barbaresca | 30 | fat-tail | Italy | 4 | [9] |
| Laticauda | 24 | fat-tail | Italy | 5 | [12] |
| Libyan Barbary | 23 | fat-tail | Algeria | 6 | [9] |
| Hammari | 7 | thin-tail | Sudan | 1,2,3 | [13] |
| Kabashi | 8 | thin-tail | Sudan | 1,2,3 | [13] |
| Sardinian | 24 | thin-tail | Italy | 4,5 | [12] |
| Comisana | 24 | thin-tail | Italy | 4,5 | [12] |
| Sidaoun | 39 | thin-tail | Algeria | 6 | This study |

### Genetic relationships amongst breeds

Pair-wise genetic relationships were estimated using identity-by-state genetic distances calculated with PLINK 1.7 [14] and graphically represented by multidimensional scaling (MDS) analysis.

### Signatures of selection analysis

To analyze genome-wide selection signatures, the MDS results were used to categorize the 23 breeds into contrasting genetic groups for comparative analysis. Table 1 summarizes the description of the breeds used in the pair-wise comparisons. The contrasting groups were as follow:

1. Ethiopian fat-tail breeds (11 breeds) vs. two thin-tail breeds from Sudan (Hammari and Kabashi);
2. Ethiopian long fat-tail breeds (6 breeds) vs. the two thin-tail breeds from Sudan;
3. Arabian peninsula fat-tail (Naimi, Najdi, Omani and Huri) vs. the two thin-tail breeds from Sudan;
4. Barbaresca vs. two Italian thin-tail breeds (Sardinian and Comisana);
5. Laticauda vs. the two Italian thin-tail breeds;
6. Libyan Barbary vs. Algerian Sidaoun.

Inter-population analyses of the six fat- vs. thin-tail groups (Table 1) were performed using the Extended Haplotype Homozygosity (EHH) – derived statistic *Rsb* [15], as in Bahbahani et al. [16–17]. To identify statistically significant SNPs under selection in each of the six pair-wise comparisons (positive *Rsb* value), one-sided *P*-values (fat- vs. thin-tail group) were derived as −logl0(1-Φ(*Rsb*)), where Φ(*Rsb*) represents the Gaussian cumulative distribution function. Interpopulation genome-wide *F_ST_* and *χ^2^* analysis were also performed to corroborate the results obtained with the *Rsb* analysis. The following constraints were introduced to define the fat-tail selection signatures: 1)−logl0 (*P*-value) ≥ 3.2, equivalent to a *P*-value of 0.0005, was used as a threshold to define significant *Rsb*; 2) candidate regions were retained if at least two SNPs, separated by ≤200 Kb, passed this threshold; 3) the candidate region was present in two or more pair-wise comparisons.

### Runs of homozygosity

Runs of homozygosity (ROHs) were estimated for each individual using PLINK 1.7 [14]. The minimum length that constituted the ROH was set to one Mb. The following criteria were also used: (i) one missing SNP was allowed in the ROH, and up to one possible heterozygous genotype; (ii) the minimum number of SNPs that constituted the ROH was set to 30; (iii) the minimum SNP density per ROH was set to one SNP every 100 kb; (iv) maximum gap between consecutive homozygous SNPs of 1000 Kb. The percentage of SNP residing within an ROH for a given breed was estimated by counting the number of times that each SNP appeared in a ROH and by dividing that number by the number of animals in each breed, allowing us to obtain a locus homozygosity range (from 0 to 1). To identify the genomic regions of “high homozygosity”, also called ROH islands, the top 0.9999 SNPs of the percentile distribution of the locus homozygosity range within each breed were selected.

### Gene annotation

Candidate regions identified by different approaches were used to annotate genes, that were either entirely or partially included within each selected region, using the NCBI Genome Data Viewer (https://www.ncbi.nlm.nih.gov/genome/gdv/browser/?context=gene&acc=10Π04604). Finally the biological functions of each annotated gene within the selection signatures was investigated via a comprehensive search of literature.

## Results

After quality control, 43,224 SNPs and 349 animals (6 to 39 per breed) were retained for the analyses (Table 1). To examine and visualize the genetic relationships among the 23 sheep breeds, we used a MDS plot of the pairwise identity-by-state distance. The results showed that most sheep breeds formed non-overlapping clusters and were clearly separate populations (Fig 1). The first dimension (C1) separated the Italian breeds from the Arabian Peninsula and African ones, likely reflecting different breeding histories. The second dimension (C2) distinguished Barbaresca from the other breeds. Therefore, with the exception of Barbaresca, the MDS grossly separated the breeds according to their genetic origin and/or to geographical proximity between their breeding areas. The Ethiopian breeds were separated into two groups: one including the three fat-rumped breeds and one short fat-tail (Molale), and one group including the long fat-tail breeds and the other short fat-tail breed (Gafera). Noteworthy, the MDS plot did not separate the breeds on the basis of the different tail phenotypes.

**Fig 1.**
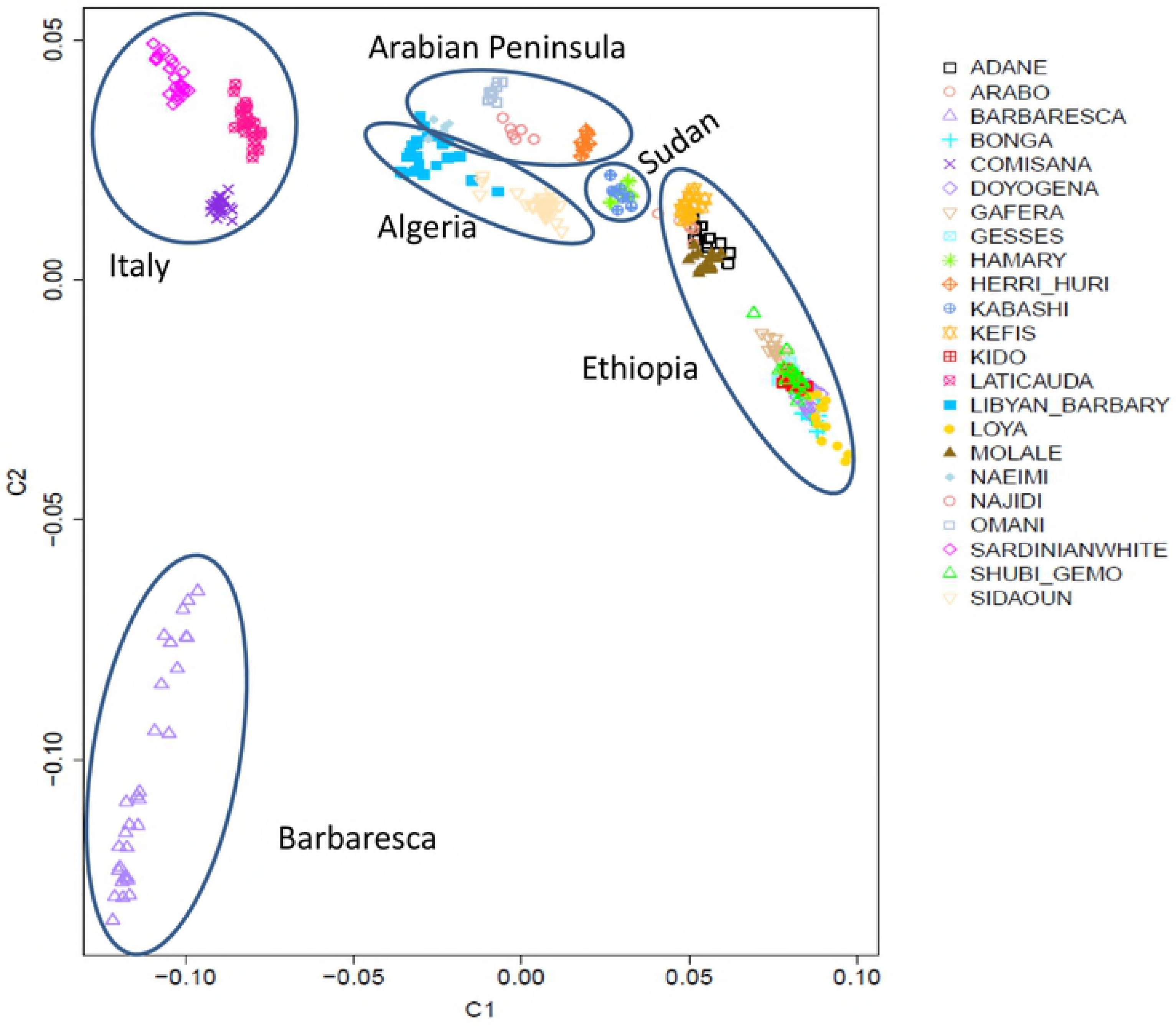
Genetic relationships among the 23 sheep breeds defined through multidimensional scaling analysis.

In this study, selection signatures for fat-tail were identified using the *Rsb* approach, after corroboration with *F_ST_* approach. The markers with a significant inter-population *Rsb* are shown in S1-S6 Tables, corresponding to the six comparisons described in the Material and methods. Likewise, in S7-S12 Tables, genome-wide *F_ST_* and *χ^2^* values, with the corresponding Bonferroni corrected *χ^2^ P*-values, are shown. A summary of the number of significant SNPs obtained with the different statistical methods in the six pair-wise comparisons is reported in Table 2.

**Table 2.** Number of significant single nucleotide polymorphisms (SNPs) obtained with the two selection signature approaches in the six pair-wise comparisons.

| Pair-wise comparison |  | No. significant SNPs |  |
| --- | --- | --- | --- |
| Fat-tail group | Thin-tail group | *Rsb*<br>P<0.0005 | Bonferroni corrected $\chi^2$ P-value $\leq 0.05$ |
| Ethiopian fat-tail | Hammary and Kabashi | 99 | 754 |
| Ethiopian long fat-tail | Hammary and Kabashi | 106 | 1,114 |
| Arabian peninsula fat-tail | Hammary and Kabashi | 64 | 108 |
| Barbaresca | Sardinian and Comisana | 638 | 2,402 |
| Laticauda | Sardinian and Comisana | 158 | 328 |
| Libyan Barbary | Sidaoun | 185 | 302 |

The genome-wide Manhattan plots where significant signals of differentiation between the fat-tail breed/group and the thin-tail of the corresponding region were shared by two or more fat-tail breed/group are reported in S1-S14 Figs. The y axis shows the probability of *Rsb* values for each marker across the genome (−log10(1-Φ(*Rsb*)). Below each plot, the positions of the significant SNPs, with their corresponding probability values, are reported for each fat-tail breed/group. *F_ST_* values and Bonferroni corrected *χ^2^ P*-values are reported only when achieved by significant SNPs located in the candidate region, or at distance ≤ 0.2 Mb up- and down-stream of the candidate region boundaries.

The majority of shared fat-tail signals were observed for the two groups of Ethiopian breeds on chromosomes (OAR) 5, 6, 10, 18 and 19 (S3, S5, S9, S13 and S14 Figs, respectively) and by the two breeds of Barbary sheep origin (Laticauda and Libyan Barbary) on OAR 3, 10, 12, 13 (S2, S9, S10 and S11 Figs, respectively). It is interesting to note that all the regions shown in S1-S14 Figs registered the presence of at least one marker attaining highly significant *Rsb* values, i.e. > 4, equivalent to a *P*-value of 0.0001. Moreover, in all the regions, except the one on OAR12 that is shared between Laticauda and Libyan Barbary (S10 Fig), at least one significant *χ^2^* value was also registered in either one or both of the fat-tail breed/groups.

Several genomic regions that frequently appeared in a ROH were identified within each breed (S15 and S16 Figs, for fat- and thin-tail sheep breeds, respectively). Table 3 provides the chromosome position, and the start and end of the ROH islands. The top 0.9999 SNPs of the percentile distribution of locus homozygosity values led to the use of different thresholds for each breed (from 0.11 to 0.83), and a total of 15 genomic regions of high-homozygosity across breeds were identified. Although the distribution of the ROH was relatively balanced and the signals were moderate in height, we found a few outstanding peaks with a high occurrence of ROH, especially in the Barbaresca breed (S15 Fig).

**Table 3.**
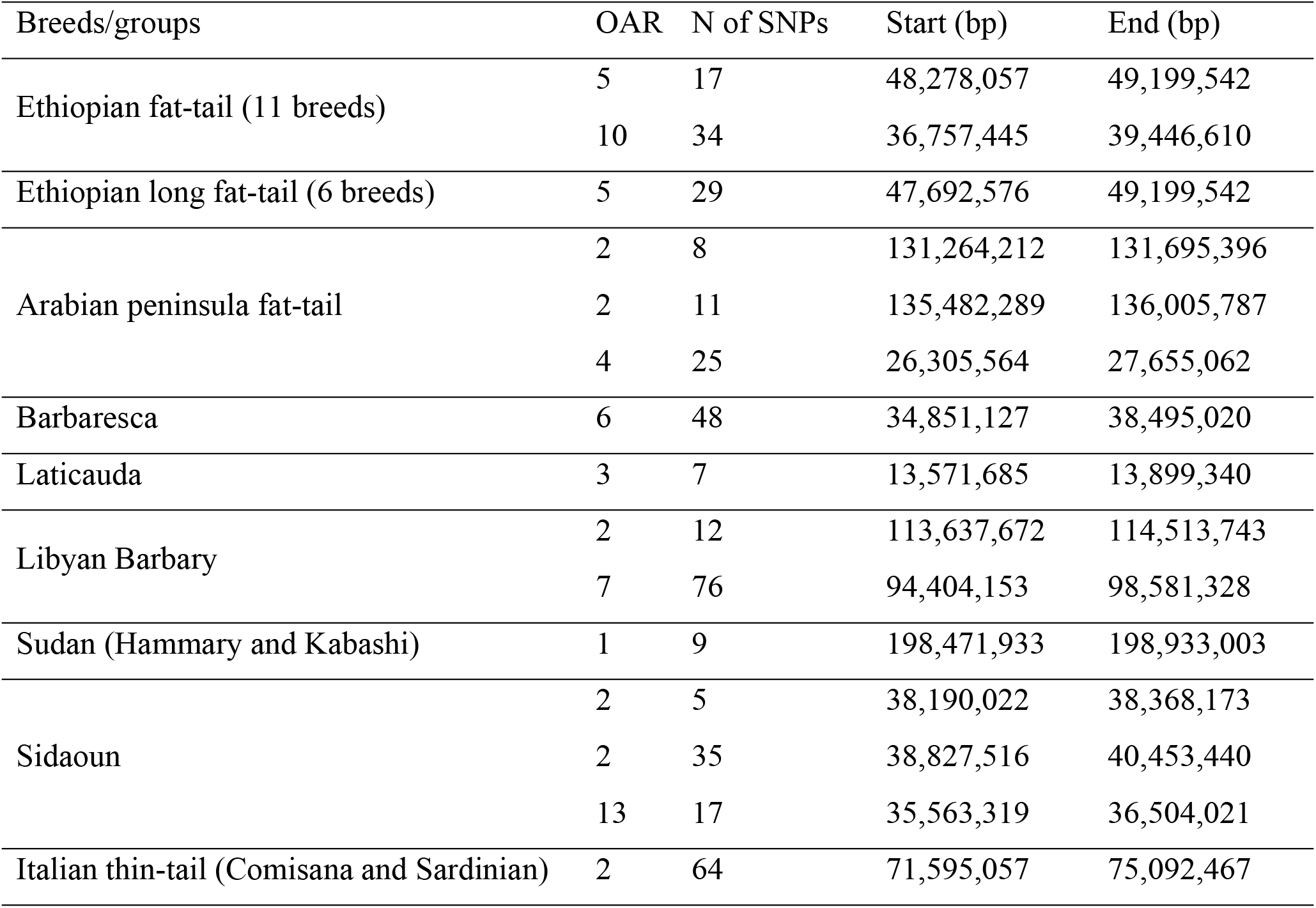
Run of homozygosity (ROH) islands identified within each breed. The chromosome (OAR), the number of single nucleotide polymorphisms (SNPs) within each ROH island and the positions of the genomic regions (in base pairs, bp) are reported.

| Breeds/groups | OAR | N of SNPs | Start (bp) | End (bp) |
| --- | --- | --- | --- | --- |
| Ethiopian fat-tail (11 breeds) | 5 | 17 | 48,278,057 | 49,199,542 |
|  | 10 | 34 | 36,757,445 | 39,446,610 |
| Ethiopian long fat-tail (6 breeds) | 5 | 29 | 47,692,576 | 49,199,542 |
| Arabian peninsula fat-tail | 2 | 8 | 131,264,212 | 131,695,396 |
|  | 2 | 11 | 135,482,289 | 136,005,787 |
|  | 4 | 25 | 26,305,564 | 27,655,062 |
| Barbaresca | 6 | 48 | 34,851,127 | 38,495,020 |
| Laticauda | 3 | 7 | 13,571,685 | 13,899,340 |
| Libyan Barbary | 2 | 12 | 113,637,672 | 114,513,743 |
|  | 7 | 76 | 94,404,153 | 98,581,328 |
| Sudan (Hammary and Kabashi) | 1 | 9 | 198,471,933 | 198,933,003 |
| Sidaoun | 2 | 5 | 38,190,022 | 38,368,173 |
|  | 2 | 35 | 38,827,516 | 40,453,440 |
|  | 13 | 17 | 35,563,319 | 36,504,021 |
| Italian thin-tail (Comisana and Sardinian) | 2 | 64 | 71,595,057 | 75,092,467 |

## Discussion

In studies aiming to detect genomic signals for specific traits, for instance signals directly associated with fat deposition and adipogenesis, the major drawback is to detect strong differences (viz. between fat-tail and thin-tail breeds) that are due either to different origins or to reproductive isolation, and not obviously involved in the trait (fat deposition). In this work, the fat-tail breeds from Ethiopia, Algeria, Arabian peninsula and southern Italy were pair-wise compared with their thin-tail counterparts from the closest geographical region. The signals that are detected between any two or more simultaneous pair-wise comparisons might consequently be considered more reliable, also because they are shown by geographically distant sheep breeds. While the fat-tail sheep of Algeria, Arabian peninsula and southern Italy were all long or semi-long fat-tail breeds, the Ethiopian fat-tail breeds included long-tail, short-tail and fat-rumped sheep breeds (Table 1). Therefore, for the Ethiopian sheep, two different pair-wise comparisons were performed, the first including all the 11 fat-tail breeds, the second including only the 6 long fat-tail breeds. The assumption here was that the first comparison might elucidate the genes involved with adipogenesis, irrespective of the shape of the tail, while the second comparison would reveal findings that are more comparable to those observed in the breeds from the other regions, and are therefore possibly more likely to be linked to the fat-tail phenotype.

We used two different, but complementary, statistical approaches to identify putative selection signatures across the phenotypically different breeds. The *F_ST_* index of differentiation is among the most widely used statistic to detect signals of selection in differentially-selected populations, where usually a locus is putatively considered under differential selection if its pair-wise *F_ST_* has a rank percentile value of 0.01 or less. Because the level of genetic differentiation between each pair of breeds/groups of the six comparisons highly varied, the decision “*a priori*” of a rank percentile to accept significant *F_ST_* may disfavor the pairs presenting the highest genetic diversity. Therefore, following Moioli et al. [12] who showed that *F_ST_* and *χ^2^* are highly correlated, in order to confirm with a statistical test which markers were significant, we calculated interpopulation locus-specific *χ^2^* values and considered significant the markers reaching a Bonferroni-adjusted *χ^2^ P*-values ≤ 0.05. Doing so, the different numbers of significant markers in each pair-wise comparison allows to appreciate their global genetic difference. However, since *F_ST_* and *χ^2^* values are based on allele frequencies and might represent an isolated event occurring by chance, and not necessarily associated with fat-tail signals, in this case, the extended haplotype homozygosity derived statistic, *Rsb*, was preferentially used. This statistic in fact considers the whole haplotype region around one marker, or group of markers, and the larger is the region in which the homozygous haplotype was maintained in the first breed in contrast with the second breed, the more reliable is the probability of carrying a fat-tail signal. The number of significant SNPs for inter-population *Rsb* is smaller than the number of significant SNPs for *χ^2^* (Table 2) confirming that the first method is more stringent. In most cases, the number of SNPs that are significant with one method showed some correlation with the number of SNPs that were obtained with the second method: the higher was the number of the significant *Rsb* values, the higher was also the number of significant *F_ST_* values. The highest number of significant signals obtained with both methods was observed in the Barbaresca breed – 1.5 % for inter-population *Rsb* and 5.6 % for *F_ST_/χ^2^*, while the lowest number was in the Arabian peninsula breeds: 0.15 and 0.25% with the two methods, respectively. In accordance with our results, Yuan et al. [11], in a selection signature analysis for tail type in sheep, identified seven and twenty-six regions using the extended haplotype homozygosity and *F_ST_* approaches, respectively, and only six small regions using both approaches. Bahbahani et al. [18] showed that the two common approaches (inter-population *Rsb* and *F_ST_)*, used to identify signatures of positive selection in East African Shorthorn Zebu, did not produce overlapping signals. Although the authors interpreted the observed absence of overlaps between *Rsb* and *FST* analyses as a possible consequence of the selection time-scale, with *Rsb* being considered more suitable for detecting signatures of recent selection, it may also be hypothesized that false positives may occur when using both *Rsb* and *F_ST_.* As the aim of this study was to identify loci most likely associated with fat-tail, we established that the candidate region(s) should be present in two or more pair-wise comparisons (Table 4). Therefore only the signatures satisfying these criteria and encompassing annotated genes will be subsequently discussed.

**Table 4.** Candidate regions and genes identified in two or more pair-wise comparisons (see material and methods). Start/end positions are based on the ovine genome sequence assembly Oar_v4.0.

| Pair-wise comparison group |  |  |  |  |  |  | Genes |
| --- | --- | --- | --- | --- | --- | --- | --- |
|  | 1 | 2 | 3 | 4 | 5 | 6 |  |
| OAR | start/end (bp) | start/end (bp) | start/end (bp) | start/end (bp) | start/end (bp) | start/end (bp) |  |
| 3 |  |  | 104,333,496<br>105,210,346 |  |  | 104,291,439<br>105,909,563 | *ZNF2, ZNF514, MRPS5, MALL, NPHP1, ACOXL, BUB1, BCL2L11, ANAPC1, MERTK* |
| 3 |  |  |  |  | 154,458,718<br>156,011,304 | 155,014,204<br>155,055,875 | *MSRB3, WIF1, LEMD3, TBC1D30, GNS, RASSF3, TBK1, SRGAP1, TMEM5* |
| 5 | 482,26,104<br>48,526,532 | 46,925,830<br>49,273,852 |  |  |  |  | *EGR1, ETF1, HSPA9, CTNNA1, SIL1, MATR3, PAIP2, SPATA24, SLC23A1, PROB1, MZB1, DNAJC18, TMEM173, ECSCR, UBE2D2, CXXC5, PSD2, NRG2, PURA, IGIP, PFDN1, HBEGF, CYSTM1, SLC4A9, TMCO6, WDR55, HARS2, NDUFA2* |
| 6 |  |  |  | 38,104,576<br>39,576,650 | 38,179,178<br>39,639,829 |  | *TRNASTOP-UCA, LOC106991224* |
| 6 | 55,697,868<br>55,794,685 | 55,697,868<br>55,811,685 |  |  |  |  | *DTHD1* |
| 6 | 75,842,854<br>76,599,033 |  |  |  | 75,533,914<br>75,941,134 |  | *ADGRL3* |
| 6 | 80,325,742<br>80,531,786 |  |  |  |  | 80,878,591<br>80,912,095 | *EPHA5* |
| 10 | 40,258,505<br>45,063,369 | 40,594,254<br>45,416,672 |  |  | 40,957,582<br>45,416,672 | 42,194,236<br>45,155,143 | *PCDH9, LOC101121526, KLHL1* |
| 12 |  |  |  |  | 43,000,381<br>43,168,807 | 43,168,807<br>43,297,179 | *RERE, SLC45A1* |
| 13 |  |  |  |  | 46,565,715<br>49,137,513 | 47,583,113<br>49,208,171 | *CDS2, PROKR2, GPCPD1, CHGB, TRMT6, CRLS1, LRRN4, FERMT1, BMP2* |
| 17 | 61,094,671<br>61,631,272 |  |  |  |  | 61,496,170<br>61,658,381 | *OAS2, OAS1, RPH3, PTPN11, RPL6, HECTD4* |
| 18 | 1,895,285<br>2,129,462 | 1,980,832<br>2,243,903 |  |  |  |  | *ATP10A* |
| 19 | 51,617,073<br>51,789,681 | 51,649,648<br>51,819,178 |  |  |  |  | *MAP4, DHX30, SMARCC1* |

Another method for detecting signatures of positive selection based on intra-population analysis is the identification of high-homozygosity regions [6]. Since ROHs are normally abundant in regions under positive selection, their accumulation at specific loci, or islands, has been used to identify genomic regions that reflect directional selection in cattle [19], sheep [20], horse [21] and goat [22]. We therefore checked if such regions of high-homozygosity overlapped with putative selection signatures in the sheep breeds considered in this study.

The region in OAR3:104.2-105.9 Mb (S1 Fig and Table 4) was identified in two comparisons: fat-tail sheep of the Arabian peninsula vs. Sudanese thin-tail sheep, and Libyan Barbary vs. Algerian Sidaoun. Out of the ten genes found in this region, only *ANAPC1* has been associated with obesity-related traits by Comuzzie et al. [23] in a Genome-Wide Association Study (GWAS) on Hispanic children. The signature on OAR3:154.0-155.6 Mb (S2 Fig and Table 4), detected in the Laticauda and the Libyan Barbary breeds, had already been reported for the Barbaresca, Laticauda and Chios breeds [9]. Yuan et al. [11], in a GWAS on seven indigenous Chinese sheep, by contrasting fat-tail versus thin-tail phenotypes, detected a signature in this region encompassing the *MSRB3*, that has been identified as a candidate gene associated with adaptation [24]. In a study on world sheep breeds, *MSRB3* was highlighted to have experienced high selection pressure [25]. However, Yuan et al. [11] warned on the possibility of distinguishing, through GWAS, false positive genes from candidate genes. On the other hand, Wei et al. [10] analyzed Chinese native sheep by contrasting thin-tail (Tibetan group) and fat-tail (Mongolian and Kazakh group) types and suggested that the genes encoded by the signal on OAR3, *i.e. MSRB3* and *LEMD3*, would be best considered as candidates for ear size. This hypothesis is supported by the fact that *MSRB3* and *LEMD3* were identified as candidate genes for ear morphology in dogs [26] and pigs [27]. Therefore, despite these genes having not been associated with the ear size [28], doubts on ascribing this signal to the fat-tail are obvious. A large region on OAR5:47.0-49.0 Mb (S3 Fig and Table 4) turned out here to encode putative fat deposition genes in the two groups of Ethiopian sheep: the one composed only by the long fat-tail, and the one including all the eleven Ethiopian fat-tail breeds. Although Fariello et al. [29] reported that this region encoded a signature differentiating prolific and non-prolific Asian sheep, the involvement of a signature in this region in the fat-tail phenotype had been previously reported [9,12]. This involvement is corroborated by genetic studies of body mass index in humans, describing a role played in obesity by the *CXXC5* gene in Americans [30] and the *PSD2* gene in the Japanese population [31]. Moreover, this selection signature overlapped with the ROH island identified in the Ethiopian fat-tail breeds, which include 17 and 29 homozygous markers respectively for the two groups of breeds (the eleven Ethiopian fat-tail, and the six Ethiopian long fat-tail). The signal on OAR6:38.1-39.6 Mb (S4 Fig and Table 4) was detected in Barbaresca and Laticauda and was previously reported as selection signature for fat-tail in these two breeds, also when compared with 13 Italian thin-tail breeds [9]. It includes the *SLIT2* gene, a potential candidate for internal organ weights in Simmental beef cattle [32] and therefore possibly connected with fat deposition. This signal is worth investigating further, because it encompasses a large region (1.5 Mb) where the Barbaresca showed 27 SNP markers with allele frequency patterns that are highly differentiated from the Italian thin-tail breeds, 15 of which exceeded the significant threshold of *Rsb P*-value < 0.0001. Moreover, this genomic region partially overlapped with the ROH island on OAR6 detected in Barbaresca and shared by more than 80% of the individuals of this breed. The Laticauda showed less significant signals in the same region, with the exception of a highly significant one (-log10(*P*-value) = 7.68) at position 38,345,613 bp. Moreover, several markers attained high *F_ST_* values and significant *χ^2^* values in the pair-wise comparisons of both breeds. However, because the two breeds have Barbary origin, it is also possible that this signature encodes loci inherited from North-African breeds. The selection signature on OAR6:55.6-55.7 Mb (S5 Fig and Table 4) was shown in this study to be present in the two groups of Ethiopian breeds; it has never been reported before as a fat-tail signal and the only gene included in the region *(DTHD1)* does not appear to have any connection with fat deposition. On the same chromosome, at position 76.5-77.5 Mb (S6 Fig and Table 4), we identified another signal in the Ethiopian group (eleven breeds) and the Laticauda, which has been reported previously in the Barbaresca but not in the Laticauda [9]. Although the thin-tail breeds analyzed in the aforementioned study for the pair-wise comparison with the Barbaresca and the Laticauda were not the same as those used here, and the methodology used to detect the signatures was also different, it is likely that this is merely a suggestive fat-tail signal. Moreover, for the only annotated gene in this region *(ADGRL3)*, to the best of our knowledge, no association with adipogenesis has been reported. Another candidate region identified on OAR6:80.6-81.0 Mb (S7 Fig and Table 4), shown by the Ethiopian group (eleven breeds) and the Libyan Barbary and including the *EPHA5* gene, was also detected in a GWAS for wool production traits in a Chinese Merino sheep population and reported to be significantly associated with fiber diameter [33]. Moreover, it has been identified through GWAS as a candidate gene for feed conversion ratio in Nellore cattle [34]. The signal on OAR7:33.5-33.9 Mb (S8 Fig and Table 4) was detected in the Barbaresca and Laticauda, as well as in a previous study [9]. While these authors could not assign an obvious role of any of the genes located in this region, Lirangi et al. [35] suggested that the *CHP1* gene is involved in cellular fat storage. Inoue et al. (2014) revealed that *OIP5* promotes proliferation of pre- and mature-adipocytes and contributes adipose hyperplasia; moreover, an increase of *OIP5* may associate with development of obesity. A common candidate region located on OAR10:40.2-45.0 Mb (S9 Fig and Table 4) observed in the two groups of Ethiopian sheep, Libyan Barbary and Laticauda, includes the *PCDH9* gene. The same signature was detected by Kim et al. [37] who contrasted the Egyptian Barki sheep with British breeds, and referred to it as signal of adaptation to arid environments. However, the Barki is a fat-tail sheep while the British are all thin-tail breeds. We would then propose the idea that this is a signal of fat-deposition, this being corroborated by the presence of *PCDH9*, reported by Wang et al. [38] as a candidate gene for obesity in humans. The signal on OAR12:43.0-43.3 Mb, shared by the Laticauda and the Libyan Barbary (S10 Fig and Table 4) has not been reported previously as a fat-tail signal and the two genes (*RERE* and *SLC45A1*) included in the region do not appear to have any connection with fat deposition. *RERE* has been identified as candidate for embryonic growth and reproductive development, whereas *SLC45A1* plays an important role in immunity related to tropical adaptation [39]. On OAR13:45.5-48.4 Mb, a selection signature shared by the Laticauda and the Libyan Barbary was identified (S11 Fig and Table 4). The region was also reported as a fat-tail signature in several studies [9–11]. The strong linkage disequilibrium between the SNPs in this OAR13 region with a missense mutation in exon 1 of the *BMP2* gene (OAR13:48,552,093-48,897,111 bp) was demonstrated by Moioli et al. [12] in the Laticauda fat-tail and Altamurana thin-tail sheep. Yuan et al. [11] emphasized that *BMP2* may play important roles in fat tail formation. However, here *BMP2* does not appear to be the only candidate gene. This signature spans a size > 3Mb, and Laticauda showed a very high *Rsb* value (-log10-Pvalue=7.96) at position 46,582,744 bp. A transcriptome profile analysis of adipose tissues from fat- and short-tailed Chinese sheep identified *CDS2* among the differentially expressed genes [40]. This gene which spans 46,560,029 to 46,605,239 bp of OAR13, encompasses the significant marker mentioned above. Another candidate region on OAR17:61.0-61.6 Mb, shared by the Ethiopian group (eleven breeds) and the Libyan Barbary (S12 Fig and Table 4) has not been detected previously as a fat-tail signal in sheep. However, Fox et al. [41] associated the *OAS2* gene found in this region to body fat distribution. Finally, the candidate regions on OAR18:1.8-2.2 Mb, and OAR19:51.6-51.8 Mb, shared by the two groups of Ethiopian breeds (S13 and S14 Figs, respectively) have not been reported previously as fat-tail signals and the five genes included in the two regions (*ATP10A* on OAR18, and *CDC25A, MAP4, DHX30, SMARCC1* on OAR19) do not appear to have any obvious connection with fat deposition. However, other studies also reported genes mapped in the same two regions with no obvious roles in sheep tail type or fat deposition [3,11]. The complexity of the fat-tail phenotype [10] may partly justify the high number of signals detected in the pair-wise comparisons.

As reported above, only two ROH islands identified in fat-tail breeds overlapped with the selection signatures. One reason for the little overlap is that ROH might detect selection related to any trait, while contrasting thin and fat tail breeds is more likely to detect signal related to this trait. However, Purfield et al. [20] reported a significant moderate correlation between the occurrence of SNPs in a ROH and two different statistical approaches (*F_ST_* and HapFLK) for identifying putative selection signatures, and showed two ROH islands that overlapped with selection signatures. Moreover, the ROH island on OAR5 identified in the Ethiopian breeds was also identified in the Lori Bakhtiari fat-tail breed [10]. The authors reported that this increase in homozygosity would be consistent with selection for mutations affecting fat-tail size several thousand years following domestication. Therefore, although the existence of ROH islands has been attributed to several factors (recombination events, demography) [42], our results corroborate the hypothesis that regions of high-homozygosity may indeed harbor targets of positive selection, as also observed in previous studies [20,22,43].

In this study, we report so far the most complete genome-wide study of selection signatures for fat-tail in sheep. We identified novel signals and confirmed the presence of selection signatures in the genomic regions that harbor candidate genes that are known to affect fat deposition. These findings also confirm the great complexity of the mechanisms underlying quantitative traits, such as the fat-tail, and further confirm the hypothesis that many different genes are involved in the phenotype. However, it is important to highlight that among the candidate genomic regions, false positives may still be a possibility. Therefore, further studies using different populations and the new ovine high-density SNP chip will be required to confirm and refine our results and investigate the role of specific genes. Notwithstanding, the selection signatures reported here provide comprehensive insights into the genetic basis underlining the fat tail phenotype in sheep.

## Data Availability Statement

This article reports no new genotyping data. All relevant data are within the paper and its Supporting Information files.

## Supporting information

**S1 Table.** Significant markers for *Rsb* in Ethiopian fat-tail breeds (eleven breeds) contrasted with two thin-tail breeds from Sudan (Hammari and Kabashi).

**S2 Table.** Significant markers for *Rsb* in Ethiopian long fat-tail breeds (six breeds) contrasted with two thin-tail breeds from Sudan (Hammari and Kabashi).

**S3 Table.** Significant markers for *Rsb* in Arabian Peninsula fat-tail breeds (Naimi, Najdi, Omani and Huri) contrasted with two thin-tail breeds from Sudan (Hammari and Kabashi).

**S4 Table.** Significant markers for *Rsb* in Barbaresca fat-tail breed contrasted with two Italian thin-tail breeds (Sardinian and Comisana).

**S5 Table.** Significant markers for *Rsb* in Laticauda fat-tail breed contrasted with two Italian thin-tail breeds (Sardinian and Comisana).

**S6 Table.** Significant markers for *Rsb* in Libyan Barbary fat-tail breed contrasted with Sidaoun thin-tail breed.

**S7 Table.** Significant markers for *F_ST_* / *χ^2^* in Ethiopian fat-tail breeds (eleven breeds) contrasted with two thin-tail breeds from Sudan (Hammari and Kabashi).

**S8 Table.** Significant markers for *F_ST_* / *χ^2^* in Ethiopian long fat-tail breeds (six breeds) contrasted with two thin-tail breeds from Sudan (Hammari and Kabashi).

**S9 Table.** Significant markers for *F_ST_* / *χ^2^* in the Arabian Peninsula fat-tail breeds (Naimi, Najdi, Omani and Huri) contrasted with two thin-tail breeds from Sudan (Hammari and Kabashi).

**S10 Table.** Significant markers for *F_ST_* / *χ^2^* in Barbaresca fat-tail breed contrasted with two Italian thin-tail breeds (Sardinian and Comisana).

**S11 Table.** Significant markers for *F_ST_* / *χ^2^* in Laticauda fat-tail breed contrasted with two Italian thin-tail breeds (Sardinian and Comisana).

**S12 Table.** Significant markers for *F_ST_* / *χ^2^* in Libyan Barbary fat-tail breed contrasted with Sidaoun thin-tail breed.

**S1 Fig.** Manhattan plot of OAR 3 depicting signals of fat-tail shared between Arabian Peninsula breeds (Naimi, Najdi, Omani and Huri) and Libyan Barbary.

**S2 Fig.** Manhattan plot of OAR 3 depicting signals of fat-tail shared between Laticauda and Libyan Barbary.

**S3 Fig.** Manhattan plot of OAR 5 depicting signals of fat-tail shared between the two groups of Ethiopian breeds.

**S4 Fig.** Manhattan plot of OAR 6 depicting signals of fat-tail shared between Barbaresca and Laticauda.

**S5 Fig.** Manhattan plot of OAR 6 depicting signals of fat-tail shared between the two groups of Ethiopian breeds.

**S6 Fig.** Manhattan plot of OAR 6 depicting signals of fat-tail shared between Ethiopian fat-tail breeds and Laticauda.

**S7 Fig.** Manhattan plot of OAR 6 depicting signals of fat-tail shared between Ethiopian fat-tail breeds and Libyan Barbary.

**S8 Fig.** Manhattan plot of OAR 7 depicting signals of fat-tail shared between Barbaresca and Laticauda.

**S9 Fig.** Manhattan plot of OAR 10 depicting signals of fat-tail shared between the two groups of Ethiopian breeds, Laticauda and Libyan Barbary.

**S10 Fig.** Manhattan plot of OAR 12 depicting signals of fat-tail shared between Laticauda and Libyan Barbary.

**S11 Fig.** Manhattan plot of OAR 13 depicting signals of fat-tail shared between Laticauda and Libyan Barbary.

**S12 Fig.** Manhattan plot of OAR 17 depicting signals of fat-tail shared between Ethiopian fat-tail breeds and Libyan barbary

**S13 Fig.** Manhattan plot of OAR 18 depicting signals of fat-tail shared between the two groups of Ethiopian breeds.

**S14 Fig.** Manhattan plot of OAR 19 depicting signals of fat-tail shared between the two groups of Ethiopian breeds.

**S15 Fig.** Regions of homozygosity in the fat-tail group/breeds. The threshold used to detect high-homozygosity regions is indicated with a black line.

**S16 Fig.** Regions of homozygosity in the thin-tail group/breeds. The threshold used to detect high-homozygosity regions is indicated with a black line.

